# Population suppression with dominant female-lethal alleles is boosted by homing gene drive

**DOI:** 10.1101/2023.12.05.570109

**Authors:** Jinyu Zhu, Jingheng Chen, Yiran Liu, Xuejiao Xu, Jackson Champer

**Affiliations:** Center for Bioinformatics, Center for Life Sciences, School of Life Sciences, Peking University, Beijing, China 100871

## Abstract

Methods to suppress pest insect populations using genetic constructs and repeated releases of male homozygotes have recently been shown to be an attractive alternative to older sterile insect technique based on radiation. Female-specific lethal alleles have substantially increased power, but still require large, sustained transgenic insect releases. Gene drive alleles bias their own inheritance to spread throughout populations, potentially allowing population suppression with a single, small-size release. However, suppression drives often suffer from efficiency issues, and the most well-studied type, homing drives, tend to spread without limit. In this study, we show that coupling female-specific lethal alleles with homing gene drive allowed substantial improvement in efficiency while still retaining the self-limiting nature (and thus confinement) of a lethal allele strategy. Using a mosquito model, we show the required releases sizes for population elimination in a variety of scenarios, including different density growth curves, with comparisons to other systems. Resistance alleles reduced the power of this method, but these could be overcome by targeting an essential gene with the drive while also providing rescue. A proof-of-principle demonstration of this system in *Drosophila melanogaster* was effective in both basing its inheritance and achieving high lethality among females that inherit the construct in the absence of antibiotic. Overall, our study shows that substantial improvements can be achieved in female-specific lethal systems for population suppression by combining them with a gene drive.

## Introduction

Sterile Insect Technique (SIT) has long been used for suppression of insect pest populations^1–4^. This method has high potential if females mate a limited number of times, so after mating with a sterile released male, the reproductive capacity of the population is reduced. However, released males often had low fitness, not just from rearing in a lab environment, but also from the methods used to sterilize the insects in the first place, usually radiation or chemicals. Use of genetic techniques to create sterile males could address some of these issues.

Release of Insects carrying a Dominant Lethal (RIDL) involves using a tetracycline-repressible lethal allele to rear insects normally in a facility. However, when released RIDL males mate with wild females, all offspring would be nonviable in the absence of tetracycline antidote. This method has two advantages. First, it does not require females to mate a limited number of times because RIDL males have viable sperm to fertilize eggs. Second, appropriately timed nonviability in the pupa stage allows RIDL offspring to still contribute to larval competition, increasing the suppressive effect of the system in species such as mosquitoes, where larval competition is intense^5^. RIDL systems have seen deployment and some success in field trials in the *Aedes aegypti* mosquitoes^6,7^, which are major vectors for dengue, zika, and other viruses. Another recent genetic SIT method involves mixing two CRISPR strains to create sterile males and eliminate females^8^. This method avoids the need for sex sorting, rearing females in release batches, and antibiotics, but this comes at the cost of increasing complexity due to the need to maintain and cross two strains. It also does not produce competitive larva, unlike the older RIDL method. The recent Engineered Genetic Incompatibility method also produces only genetically sterile males, having the advantage of using only one strain but still requiring antibiotics^9^.

Female-specific Release of Insects carrying a Dominant Lethal (fsRIDL) uses sex-specific splicing to avoid lethality in males. This allows sex sorting and reduces the need for antibiotics in the rearing facility because insects scheduled for release are reared without antibiotics, which also eliminates females. fsRIDL alleles can also persist in the population for multiple generations, only being removed when they cause lethality in females. This significantly increases the suppressive power of the system, reducing the necessary size of male releases. Developed in the crop pests *Ceratitis capitata*^10^ and *Drosophila suzukii*^11^, fsRIDL has also already undergone field trials in the mosquito *Aedes aegypti*^12^.

All of these techniques are specific to the target species and are also self-limiting. However, even in the latest fsRIDL systems, necessary release sizes tend to be very large, and repeated releases are needed to eliminate a population and potentially to keep new migrants from re-establishing the target population. This limits the application of fsRIDL and similar genetic control methods.

Engineered gene drives may provide a solution because they can spread autonomously from a small, single release. In homing suppression drives, which have been demonstrated in flies and mosquitoes in the laboratory^13,14^, the drive allele usually uses CRISPR/Cas9 to cleave a haplosufficient but essential female fertility gene in the germline. The break undergoes homology-directed repair, copying the drive allele and thus enabling more offspring to inherit the drive. Eventually, the drive will create many sterile female homozygotes when it reaches high frequency. If end-joining repair occurs, the guide RNA (gRNA) may no longer be a match for the new target sequence, which is termed a resistance allele. If these alleles preserve the function of the target gene, population suppression will fail. However, these can be avoided by using multiplexed gRNAs and targeting conserved sites, which makes resistance alleles nonfunctional with lower fitness. Modification drives can similarly eliminate nonfunctional resistance alleles by targeting essential target genes and providing a rescue^15,16^. However, nonfunctional resistance alleles can still seriously impact suppression drives. They can substantially reduce the power of the drive, which is also negatively affected by somatic Cas9 expression and imperfect drive conversion^14,17^. For many drives, this can prevent successful population suppression, especially in complex spatial environments^18–21^. Another disadvantage of homing suppression drives is that they tend to spread without limit unless specific measures are implemented, which are still being developed and may have varying effectiveness depending on the size of the target population.

By combining fsRIDL and homing gene drive, it may be possible to mitigate some of the disadvantages of each system. This was initially proposed in the earliest RIDL study, assuming perfect drive inheritance^22^, and a similar system was modeled later^23^. By allowing drive heterozygous males to pass on the fsRIDL allele at a high rate, such alleles will persist in the population longer, and so the system would have a stronger suppressive effect for the same release size. However, the performance of these systems was not comprehensively modeled (the drive was assumed to be ideal^22,23^), nor has such a system demonstrated in insects.

Here, we analyzed the effect of several performance parameters on a combined homing drive-fsRIDL system, which we call Release of Insects with Dominant Drive female-Lethal Effectors (RIDDLE). We assessed the effect of drive conversion, fitness costs, resistance allele formation, and incomplete lethality in both general and mosquito-specific models, finding that even modest level of drive conversion can substantially increase the suppressive power of the system. We also constructed a RIDDLE system in *Drosophila melanogaster* and demonstrated that it functioned successfully for both gene drive and female lethality in the absence of antibiotic. Together, these results show that RIDDLE is a potentially promising tool for suppression of disease vectors, agricultural pests, and invasive species.

## Methods

### Population suppression strategies

We simulated fsRIDL by making all females with the construct nonviable. Homozygous males are still released because an antibiotic is used in a rearing facility to prevent female mortality. We simulated SIT by using sterile males, which prevent females from reproducing if they mate with them.

In the Release of Insects with Dominant Drive female-Lethal Effectors (RIDDLE) system, a female-specific dominant lethal gene is the cargo of a homing drive. Female individuals carrying one or two such drive alleles are nonviable (Figure 1). The drive can convert the wild-type allele to another drive allele in germline cells through Cas9-mediated cleavage and homology-directed repair with a probability equal to the *drive conversion rate* (the drive efficiency).

**Figure 1.**
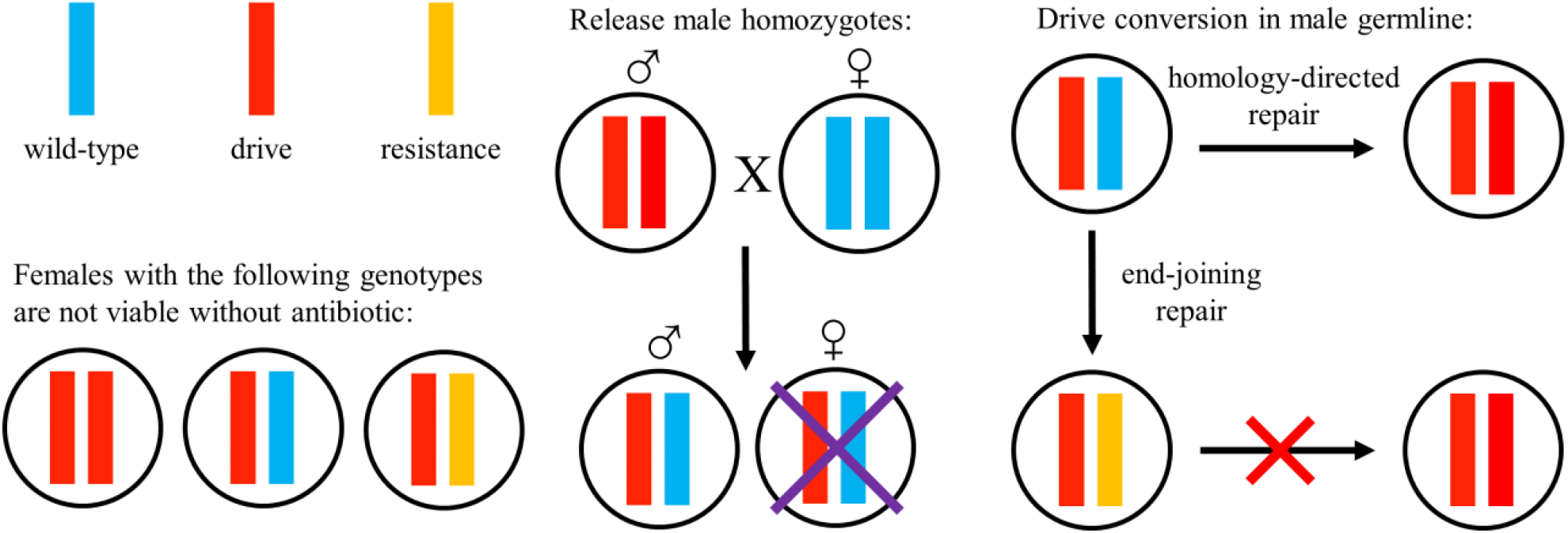
The RIDDLE system. Females with one or more drive alleles are nonviable. Male homozygotes are released into a population, and their daughters will be nonviable. Drive conversion can take place in male germline cells, allowing over half of the progeny to inherit the drive. A resistance allele can form as an alternative to successful drive conversion, and such alleles cannot be converted into drive alleles.

Also, the wild-type allele may become a resistance allele through end-joining repair with a chance equal to the *germline resistance formation rate*. Unlike a wild-type allele, a resistance allele cannot be converted to a drive allele, since end-joining repair usually changes the sequence near the cut site (Figure 1). Resistance alleles potentially have selection advantage compare to the drive because they do not make females nonviable. We initially modeled a scenario where resistance alleles were fully viable (“r1” alleles that fail to disrupt any target gene). However, if the drive targets an essential gene in a conserved region with multiple gRNAs, nearly all resistance alleles will disrupt the function of the gene (Figure S1). These will be nonfunctional (or “r2”) resistance alleles. If the target gene is haplosufficient, then resistance allele homozygotes will be nonviable (in both males and females). Similarly, if the target gene is haplolethal, all individuals with even one resistance allele will be nonviable.

### Discrete generation model

The discrete generation panmictic model simulates a random-mating population with non**-**overlapping generations using the genetic simulation framework SLiM (version 4.0.1)^24^ (Figure S2). For a full list of default parameters, see Table S1. In this model, each female randomly selects a male for potential mating, and the chance of actual mating is proportional to the male’s *genotype-based fitness*. If the female does not mate, she can select another male, up to a maximum of 20 attempts. Wild-type fitness is set to 1, and the net fitness of an individuals is the multiple of the fitness of all of their alleles. To characterize density-dependent competition, we assume that only females contribute to competition, indirectly representing competition between offspring (due to large releases of males, the ecology would be drastically changed if males contributed to competition, so we do not model this). The fecundity of a female is:

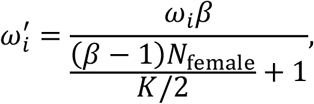

where *ω*_*i*_ is her genotype-based fitness, β is the low-density growth rate (representing the growth rate when the population is close to zero), *K* is the carrying capacity of the population, and *N*_*female*_ is the female population size. The number of offspring from a single female is selected from a binomial distribution with 50 trials and a probability of 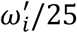. After the reproduction stage, the parental generation is removed. Female drive carriers are not removed from the population instantly, but will be counted in the population size and thus contribute to density-dependent competition. However, they are sterile, representing early mortality.

The *drop ratio* is defined as the ratio of release size of male homozygotes to half the carrying capacity (representing the normal number of males in the population), and this number is released every generation. The simulation end when the population is eliminated or after 100 generations of male releases.

### Mosquito model

We also used a previous developed mosquito model^21,25^ that simulates various life stages, including larval competition (Figure S3). For a full list of default parameters, see Table S2. In this model, a female mosquito may live up to 8 weeks, while a male mosquito may live up to 5 weeks. Individuals are in the juvenile/larvae stage during the first two weeks in the life cycle and will enter adulthood in their third week. Adult females may mate multiple times during their lives, but have only a 5% weekly chance to remate if they have already mated (otherwise they will reproduce using the original mate’s sperm). A mated female has a 50% chance to reproduce (which can take place in 3-week old females), and the number of her offspring is drawn from a Poisson distribution with a mean of 50.

Because adult mosquitoes do not have strong direct competition, they have density-independent survival rates,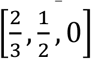 for males and 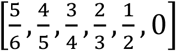 for females. Juveniles, however, face density-dependent competition and have a reduced survival rate in their first week, which can be calculated as follows:

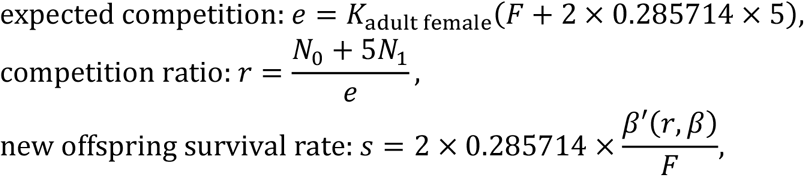

where

- *K*_*adult female*_ is the population size of adult females at the environment’s carrying capacity,
- *F* is the expected number of offspring per adult female (25),
- *N*_0_ and *N*_1_ are the population sizes of new juveniles (week 0) and older juveniles (week 1), respectively,
- 5 indicates that the older larvae contribute five times as much competition as new juveniles,
- 0.285714 indicates the relative population size of older female juveniles compared to the adult female population size,
- *β*’(*r, β*) is the relative growth rate, a function of the competition ratio *r* and the low-density growth rate *β*. In our models, β is a constant, and *β*’ is a monotonic descent function of *r* due to competition (Allee effects are not simulated in our panmictic model). When *r* → _0_, *β*^’^ (*r*) → *β*. The specific form represents the *density-dependent growth curve* of the population. This curve can be concave, linear or convex (Table S3, Figure S4), demonstrating different resource utilization strategies of the mosquito population^26,27^.

A certain number of adult male drive homozygotes are released into the population every week (all three weeks old, so released within one week of becoming adults). The release size is governed by the *drop ratio*, defined as ratio of the number of released individuals per generation (∼3.167 weeks) to the number of adult males when the population is at is normal capacity:

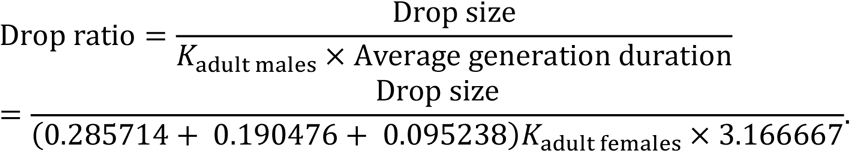

Female drive carriers take part in the density-dependent competition during their juvenile/larvae stage, but will be removed at the end of the second week before becoming adults, representing a delayed lethal effect^5^. Individuals that are nonviable because of nonfunctional resistance alleles are immediately removed from the population without contributing to competition. Simulations stop when the population is eliminated or after 317 weeks (∼100 generations).

### Data generation and analysi

SLiM simulations were performed on Polaris HPC at Peking University. Data analysis was performed in Bash and Python, and figures were prepared using Python and MATLAB. All SLiM scripts and data are available on GitHub (https://github.com/jyzhu-pointless/RIDL-drive-project/). An interactive demonstration of the models can be found online (https://jyzhu-pointless.github.io/Gene-drive-playground/#riddle).

### Plasmids construction

The tTAV with tetO element DNA was synthesized by the company BGI, and the drive plasmid was built with restriction digestion, PCR, and HiFi assembly. After transformation in DH5α competent *Escherichia coli*, plasmids were purified with ZymoPure Midiprep kit for microinjection and Sanger. The plasmid was confirmed by Sanger sequencing. Plasmid sequences are available at GitHub (https://github.com/jchamper/ChamperLab/tree/main/Drive-RIDL).

### Generation of transgenic lines

Microinjection was conducted by the company UniHuaii. The donor plasmid (500 ng/µL), along with the Cas9 helper plasmid TTChsp70c9^28^ (500 ng/µL) and gRNA helper plasmid XX22-TTTrgU2t (100 ng/µL), was injected into *Drosophila melanogaster* flies containing the haplolethal homing drive AHDr352v2^15^. After hatching, injected G0 individuals were crossed with the *w*^*1118*^ in vials with tetracycline to produce transgenic offspring. Successful transformants were marked with EGFP, indicating the presence of the new drive.

### Fly maintenance

Flies were housed in an incubator at 25°C and 60% relative humidity with a 14/10-hour day/night cycle. To prevent the death of transgenic female offspring, we added 500 µl of 1 g/L tetracycline solution to food vials after the food had solidified. Vials were left at room temperature for at least 72 hours before adding flies to ensure that the tetracycline solution was fully absorbed by the food.

### Drive performance

The RIDDLE line was crossed with a *nanos*-Cas9 line marked by DsRed to generate drive offspring with both DsRed and EGFP fluorescence. These drive offspring were further crossed to *w*^*1118*^ flies, and their progeny were screened for fluorescence. The percentage of EGFP flies represented the drive inheritance rate. In egg viability experiments, individual females were given 20-24 hours to lay eggs for several days in a row and were transferred to a new food vial each day. The offspring were later phenotyped up to 17 days post oviposition, to ensure that only progeny of the original females were counted.

## Results

### Comparative suppression performance in the discrete generation model

Using our discrete-generation model, we sought to compare the performance of the Release of Insects with Dominant Drive female-Lethal Effectors (RIDDLE) system to fsRIDL and SIT (Figure 2). We found that SIT had the worst performance as expected, requiring relative release ratios above 0.46 (referring to the number of males dropped compared to the normal number of males in the population at equilibrium before any population suppression) to eliminate the population. fsRIDL (equivalent to RIDDLE with a drive conversion efficiency of 0) required only a release ratio of 0.24 because female offspring can still provide competition, and male offspring can still pass on the fsRIDL allele to half of their own offspring. For RIDDLE, a greatly increased success range can be seen as drive conversion efficiency increases, and indeed, with the drive was nearly ideal, even the smallest tested release size of 0.03 was successful in eventually eliminating the population.

**Figure 2.**
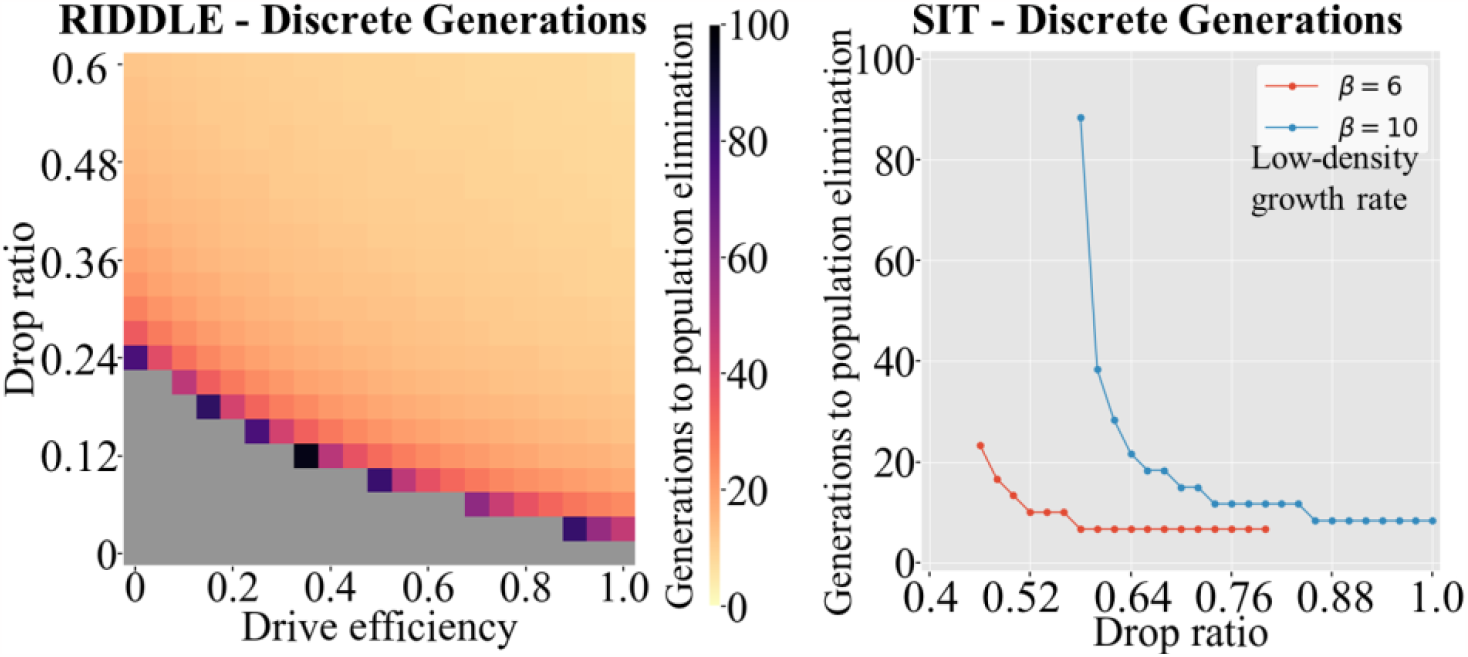
RIDDLE performance in the discrete-generation model. Males homozygous for RIDDLE (left - equivalent to fsRIDL when drive efficiency is zero) or SIT males (right) were released into a population every generation at the drop ratio, the relative number compared to the normal male population at equilibrium. Both graphs show the average time to population elimination based on 20 simulations. The low-density growth rate for the RIDDLE allele is 6. Grey indicates failure to eliminate the population after 100 generations.

All of these release sizes were small for two reasons. First, the release ratio only represents the “effective” release in our model, assuming released individuals are equivalent to wild-borne males. In reality, this is not likely to be the case, with released males having generally worse mating performance. This factor would affect all three drive systems equally, assuming that SIT was based on genetic methods and not radiation or chemical-sterilization, which would tend to have higher fitness costs. Second, the model itself is highly amenable to population suppression by continuous releases. This may be because in the model, competition affects rather than offspring viability, but also because the density growth response is a “concave”-shaped Beverton-Holt curve, which allows for stability if the population is perturbed in discrete-generation models, but also results in substantial population reduction if there is only modest suppressive power. While not necessarily a major issue for standard gene drive performance assessment, the repeated release nature of RIDDLE means that the drive has a major advantage if the population can be initially strongly suppressed with modest effort, because the effective release ratio of males compared to the number of males in the actual population will then greatly increase, further increasing the suppressive power of the system.

### RIDDLE performance in a mosquito model

To assess our drives in a more realistic setting, we used a mosquito model with specific life stages and competition only at the juvenile/larvae stage. We also adapted this model to have three different density response curves, the “concave” Beverton-Holt curve, a linear curve, and a “convex” curve (Figure S4). With a low-density growth rate of 10, the concave curve in the mosquito model needed a substantially higher minimum release size (2.6) to achieve population elimination because competition was now only between juveniles (Figure 3). For the linear density response curve, the release ratio was far greater (14 for fsRIDL), with a similar increase for convex curve (25 for fsRIDL). A similar pattern was seen for SIT (Figure S5). Population elimination in these other curves was far less effective was because higher suppression power was needed to initially reduce the population and further increase the actual release ratio. However, in all cases, higher drive conversion efficiency allowed for far smaller releases sizes, and indeed, the reduction in the release size was directly proportional to the drive conversion efficiency.

**Figure 3.**
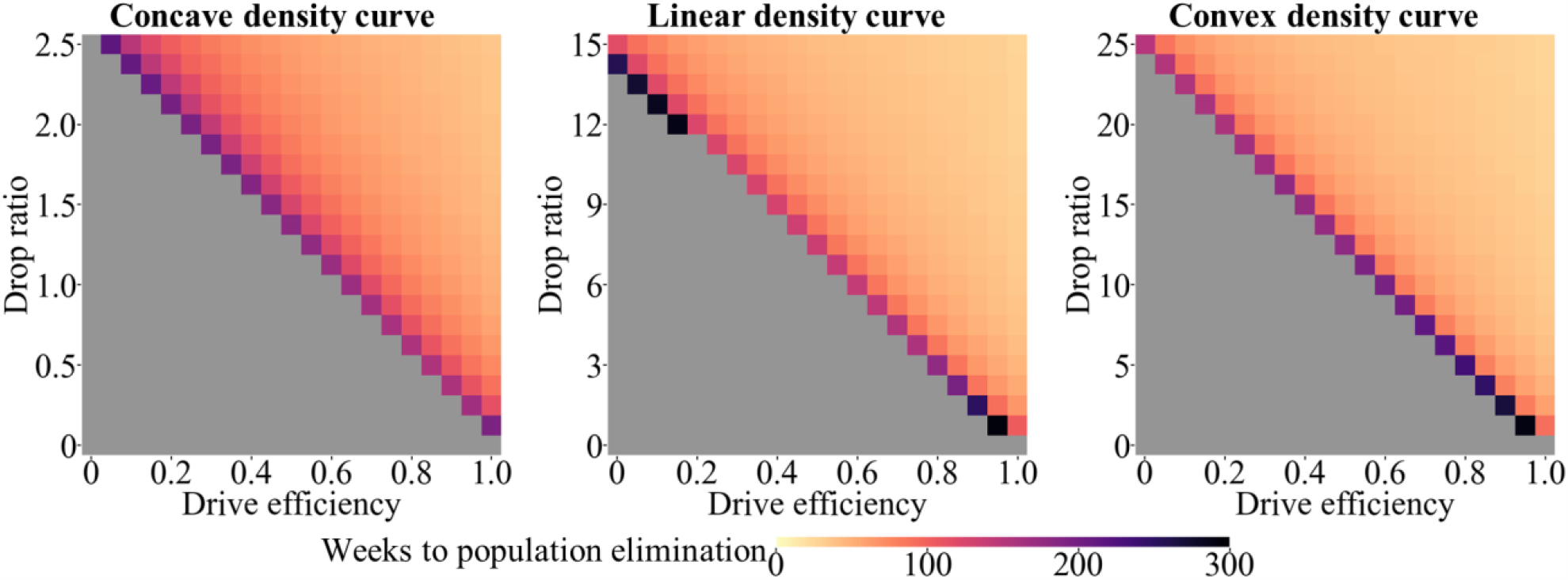
Effect of density dependence in the mosquito model. Males homozygous for RIDDLE were released into a population every week based the drop ratio, which specifies the relative number released each generation (3.17 weeks) compared to the male population at equilibrium. The density dependence of the model was varied (see Figure S4), with a fixed low-density growth rate of 10. Grey indicates failure to eliminate the population after 100 generations (317 weeks).

We also assessed the effect of varying the low-density growth rate parameter on our suppression drive constructs, using a linear density response curve (Figure S6). We found that the necessary release ratio was proportional to the low-density growth rate, as expected. Considering the major potential seasonal and ecological^25^ variation in this parameter, it is thus a critical consideration for developing a successful release program.

While most gene drives do not have substantial unintended fitness costs aside from possible cargo genes or somatic targeting of essential genes in suppression drives, some fitness costs are occasionally observed. Using a linear density response and a low-density growth rate of 6, we thus investigated the effect of fitness costs of RIDDLE, fsRIDL, and SIT. We found that unless males had very low fitness, release sizes would not be substantially affected (Figure S7). Note that this refers to genetic fitness cost affecting all drive carriers, not to any additional fitness costs of reared insects compared to wild-borne insects.

### Effect of resistance alleles on RIDDLE suppression power

All CRISPR-based gene drives are potentially vulnerable to resistance allele formation, and RIDDLE is no exception. Resistance alleles would allow females to remain viable. However, unlike homing suppression drive, where a single functional resistance allele (referring to alleles that preserve the function of the drive’s target gene) could prevent suppression, resistance alleles for RIDDLE would merely reduce its power. These could be “functional” if there is a target gene, or could refer to any resistance allele if there is no target gene and all resistance alleles have the same fitness effects. If the entire population was composed of such alleles, then RIDDLE would act identically to fsRIDL, still retaining moderate suppressive power. To investigate the effect of resistance, we fixed the drive conversion efficiency to 50% and implemented a germline resistance allele formation rate, allowing some of the wild-type alleles to be converted to resistance alleles (Figure 4). We found that when the germline resistance allele formation rate was lower than around 10^−4^, the necessary release ratio to suppress the population was largely unchanged from a scenario without resistance alleles (which required a male release ratio of 3 to achieve suppression). When resistance allele formation was maximum (0.5 - so no remaining wild-type alleles after drive effects in male drive heterozygotes), RIDDLE was substantially impaired, and the required release size was slightly below 5. This was still a small improvement over fsRIDL (which required a release ratio of 6 under similar conditions).

**Figure 4.**
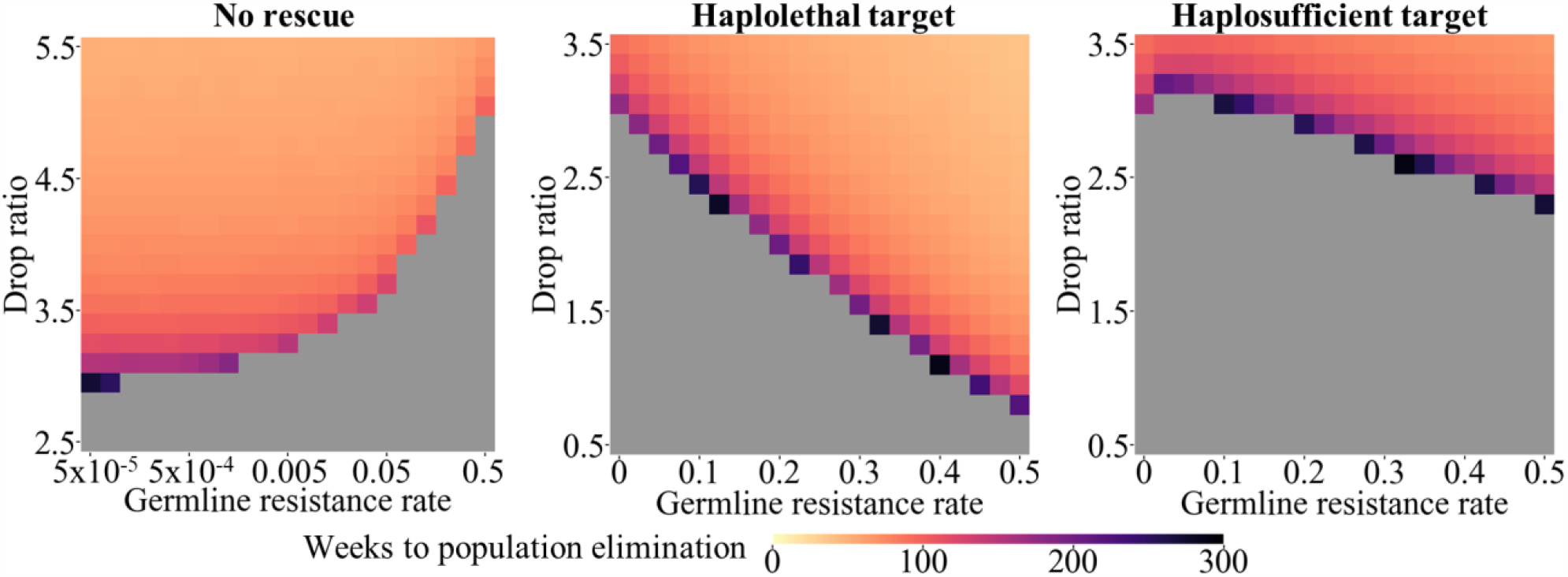
Effect of resistance alleles in the mosquito model. Males homozygous for RIDDLE were released into a population every week based the drop ratio, which specifies the relative number released each generation compared to the male population at equilibrium. With a low-density growth rate of 6 and a drive conversion efficiency of 50%, the resistance allele formation rate was allowed to vary. The left panel assumes all resistance alleles are functional (note the logarithmic scale), while the center and right panel assume nonfunctional resistance alleles. Grey indicates failure to eliminate the population after 100 generations (317 weeks).

Homing modification drives have been developed that target an essential gene and provide rescue, thus allowing nonfunctional resistance alleles to be removed^15,16^. Use of multiplexed gRNAs and conserved target sites could allow nearly all resistance alleles to be nonfunctional^29^. We thus modeled two rescue strategies where all resistance alleles are nonfunctional, with RIDDLE drives that target with haplolethal genes or haplosufficient but essential genes (Figure 4). We found that nonfunctional resistance was actually substantially beneficial to the haplolethal targeting drive. This is because any individuals inheriting a resistance allele are immediately nonviable. Such alleles thus contribute to population suppression, even if not as much as drive alleles. The RIDDLE allele with a haplosufficient target saw a similar effect, but greatly reduced in magnitude because only resistance allele homozygotes are nonviable. Indeed, a small resistance allele formation rate actually very marginally hampered this drive because such resistance alleles can prevent drive conversion in male drive carriers. These results suggest that a RIDDLE drive using Toxin-Antidote Recessive Embryo (TARE) as the driving element in lieu of a homing drive could also improve suppressive power. Modeling indicated a substantial improvement compared to fsRIDL if the total germline cut rate was 100%, though not quite as much as a homing drive (Figure S7).

### Demonstration of RIDDLE in flies

To demonstrate the concept of RIDDLE in a model organism, we modified our haplolethal homing rescue drive to contain a tTAV element with a tetO/hsp70 promoter (Figure 5A). This allows the tTAV (which binds to tetO to increase expression) to have runaway lethal expression in the absence of tetracycline, but only in females due to the presence of a female-specific intron.

**Figure 5.**
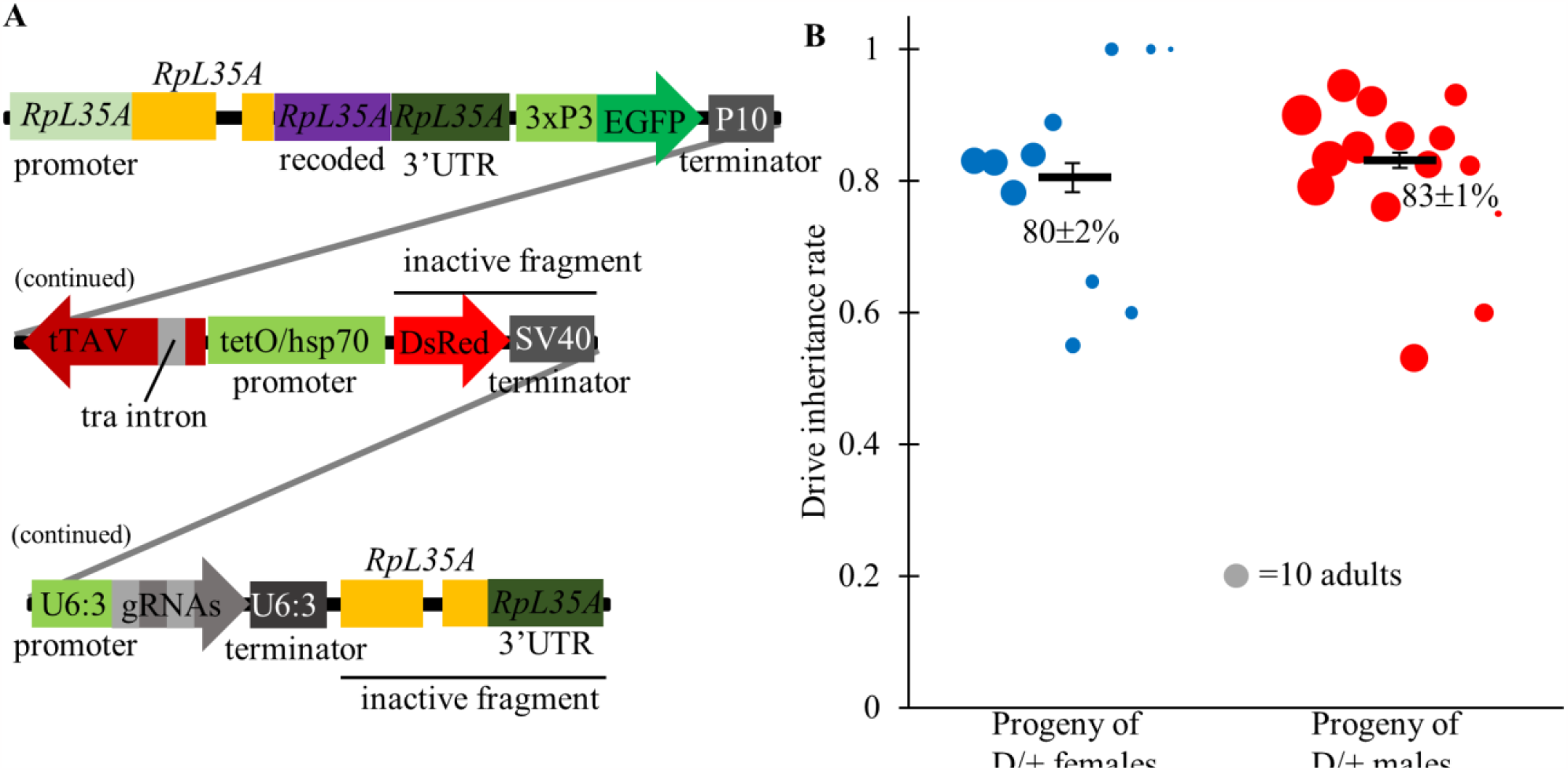
Schematic of RIDDLE and homing drive performance. (**A**) The RIDDLE allele is transformed into a previous split homing rescue drive, rendering the DsRed marker inactive. The RIDDLE allele has an EGFP gene expressed in the eyes, and a tTAV gene that can activate itself in the absence of tetracycline. The drive retains a recoded rescue copy of its haplolethal target and two gRNAs. (**B**) The drive showed high inheritance from male and female heterozygote parents (with one paternal copy of Cas9) in the presence of tetracycline. Each dot represents progeny from a single drive individual. Black bars represent the average and standard error.

To assess drive performance, the RIDDLE line was crossed to a *nanos*-Cas9 line (Cas9 with the *nanos* promoter, 5’ UTR, and 3’ UTR) in the presence of tetracycline to generate drive offspring with both DsRed and EGFP fluorescence. These drive offspring were further crossed to a *w*^*1118*^ line, and their progeny were screened for fluorescence. The percentage of EGFP flies represented the drive inheritance rate, which was 80% for female heterozygotes and 83% for male heterozygotes (Figure 5B, Data Set S1). Though moderately high, this was somewhat lower than the 90% of the haplolethal homing drive without tTAV^15^, perhaps due to the larger size of the RIDDLE construct.

To assess the ability of the RIDDLE allele to induce female nonviability, several crosses were conducted, all in a Cas9 homozygous background. First, control crosses with only Cas9 revealed a high egg-to-adult viability of 93% with tetracycline and 85% without tetracycline (Figure S8, Data Set S2). Tetracycline is not likely to have a large effect on viability, so this difference was perhaps due to batch effects of the flies or from the food. To assess RIDDLE heterozygote viability, most representative of field conditions, RIDDLE drive homozygous males were crossed to Cas9 females. All offspring were thus heterozygous. Viability was 95% with tetracycline and 56% without tetracycline (Figure 6A-B, Data Set S2). Without tetracycline, 76% of progeny were males, indicating that females had 31% relative viability. This high viability may be due to genomic position of the construct, or it could be because the tTAV gene relies on the P10 terminator that is oriented for the EGFP gene, possibility leading to reduced tTAV mRNA stability. We also performed a cross of RIDDLE drive heterozygous males and Cas9 females in the absence of tetracycline (Figure 6C, Data Set S2). Here, the drive inheritance remained at 83% and the female drive carrier relative viability at 33%, indicating that drive and female lethality could occur together. Unusually, non-drive progeny had a high female bias (Data Set S2), though the reasons for this were unclear. Finally, we assessed drive homozygotes as would be found a rearing facility. Viability was lower, even in the tetracycline vials, likely due to poor food batches (Figure 6D, Data Set S2). However, in the absence of tetracycline, no females were viable, indicating an efficient construct (Figure 6E, Data Set S2).

**Figure 6.**
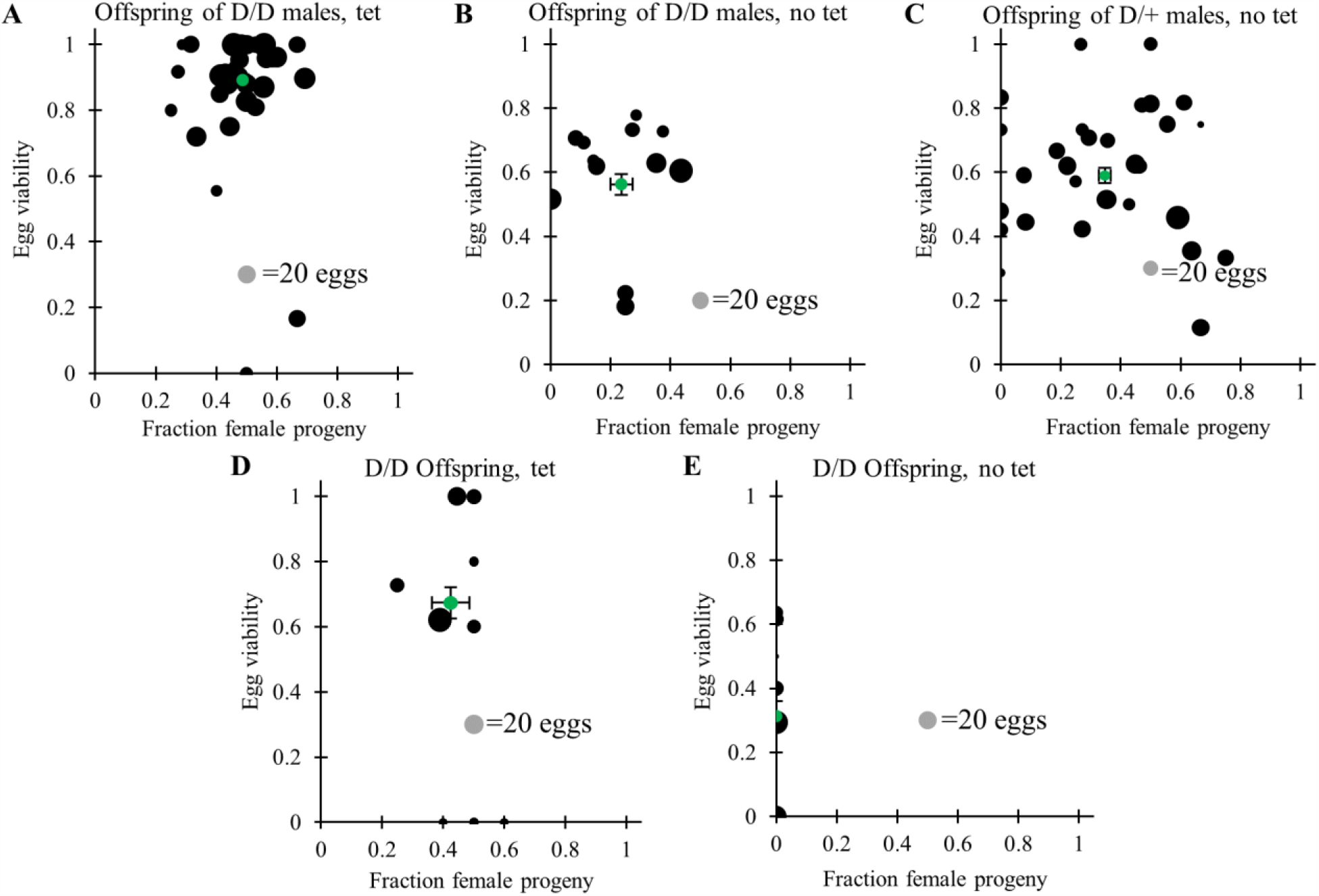
Female lethal effect of the RIDDLE allele. Egg viability was recorded in vials in various crosses, all of which were with a Cas9 homozygous background. Adult progeny were also phenotyped for sex. Crosses were between drive homozygotes males and wild-type females (**A**) with and (**B**) without tetracycline, (**C)** between drive heterozygous males and wild-type type females without tetracyclines, and between drive homozygous males and females (**D**) with and (**E**) without tetracycline. Each dot represents progeny from a single drive individual. The green dot represents the mean for all individuals, and black bars represent the average and standard error.

Because some female heterozygotes survived, we were curious to check their relative fitness compared to non-drive females. However, these females appeared fertile and did not have significantly different fecundity than Cas9 females (Data Set S3). It is possible that they may suffer other fitness costs such as reduced longevity, but these were not assessed. For RIDDLE, the high female viability would be close, but not quite enough for the drive to persist in a population (as opposed to being self-limiting due to female-lethal effects), based on drive performance and estimated embryo resistance allele formation rate in females^15^, but only if the drive was a “complete” drive rather than a split drive, which would always be self-limiting.

Because incomplete lethality could potentially affect release candidates, we modeled this with a low-density growth rate of 6, linear density response, 50% drive conversion efficiency, and 50% resistance allele formation rate. RIDDLE homozygous females still had zero viability. For both haplolethal and haplosufficient targets, high female survival actually resulted in more efficient population elimination because the drive gained some character of a self-sustaining homing suppression drive (Figure S9). This could potentially allow RIDDLE strategies to succeed even in species where a nearly perfect RIDL element could not be constructed. However, for the haplosufficient target, female survival over 30% prevented successful population suppression with any release size because the resistance alleles inhibited drive conversion, essential to produce nonviable female homozygotes. With lower resistance allele formation rates, the haplolethal target drive would also suffer from similar effects if drive conversion remained low. Depending on the drive conversion rate and fitness costs, sufficiently high female heterozygote viability could also prevent the RIDDLE system from being self-limiting.

## Discussion

In this study, we investigated the possibly of combining homing drive and female-specific RIDL strategies. Building on previous work^22,23^, we extensively modeled fsRIDL-drive (RIDDLE) and found that it was substantially more efficient than fsRIDL alone, which was itself a major advancement of RIDL and SIT methods. This high efficiency, still depends on RIDDLE’s ability to maintain high drive conversion while either maintaining low resistance allele formation or targeting essential genes and ensuring that resistance alleles are nonfunctional. Our RIDDLE demonstration in *Drosophila melanogaster* serves as a proof-of-principle, having effective homing and RIDL elements.

In principle, if the drive conversion is perfect in the absence of fitness costs, even arbitrarily small but continuous releases of a RIDDLE system could eventually suppress a population. More practically, an intermediate level of drive conversion would greatly reduce the necessary release effort compared to fsRIDL, generally reducing the required release size proportionally to the drive conversion rate. The release size can be somewhat further reduced if the drive targets an essential gene and produces moderate amounts of nonfunctional resistance alleles, with this effect being substantially larger if the resistance alleles are haplolethal. Similarly, a RIDDLE drive based on TARE^30–32^ instead of homing drives for the driving element also provided considerable improvement compared to fsRIDL, albeit less than a homing drive. This could also potentially allow a RIDDLE strategy in haplodiploid organisms (where homing drive can only function in females^33,34^) to provide some increased efficiency compared to fsRIDL, as well as other organisms where high CRISPR cut rates can be achieved (for efficient TARE), but not high drive conversion rates.

Basing the driving element on TARE would also provide an increased measure of safety for the drive in terms of confinement. While RIDDLE is intrinsically a self-limiting drive system, random mutations that inactivate the female-lethal cassette could form a complete drive system. Such mutations would be more common during homology-directed repair (necessary for drive conversion) than for normal DNA replication. Because the drive is dominant female-lethal, such alleles may not escape elimination from repeated releases, but it might be possible for individuals with such mutations to migrate out of the release area. This could cause the drive to spread in an unconfined manner, which would be heavily mitigated if the drive was a more confined TARE system. Another way to address this would be use of a split drive in which the Cas9 is on a separate genomic locus and thus not copied by drive conversion. Though this would sacrifice some performance, the repeated release method of RIDDLE would mean a much smaller loss in overall drive effectiveness compared to the loss experienced in a standard single-release gene drive strategy when switching to a split drive system. Daisy chains^35,36^ and closely linked Cas9 alleles could further mitigate this loss of effectiveness while still ensuring the drive would be confined if the female-lethal element becomes mutated.

Another way for RIDDLE to lose confinement is for the female-lethal element to have less than 100% lethality before the females can reproduce. We observed this in our experimental demonstration, with female heterozygotes having 30% survival compared to males and still retaining the ability to reproduce effectively. Female homozygotes were still completely nonviable in the absence of tetracycline. If heterozygote survival is sufficiently high, the drive may function more like a standard female fertility suppression gene drive, spreading widely, but still suppressing the population if drive conversion efficiency is high enough. If survival is too high, then the suppressive power of the drive may be lost if drive conversion is modest. Avoiding this could involve alternate stronger tTAV prompters or utilizing genomic insertion sites that allow for higher transgene expression. However, we found that small levels of female heterozygote survival actually benefit the suppressive power of the system because such females can contribute to RIDDLE allele spread.

Similar systems to RIDDLE exist, such as homing suppression drives with high somatic expression that renders females sterile. Both constructs would have similar principles, but RIDDLE would likely be more efficient unless total germline cut rates are high (drive conversion need not be high in these cases) because homing drives would need to be released as heterozygotes, and it would be undesirable to add viable wild-type alleles to the population. If homing suppression drives can make dominant-sterile resistance alleles^37^, a repeated release system based on them would be even more efficient than RIDDLE. However, RIDDLE would still have the advantage of easier rearing because female homozygotes are viable in the presence of antibiotic, preventing the need to cross drive males to wild-type females for line maintenance.

With fsRIDL now demonstrated in three different pest species (*Ceratitis capitata*^10^, *Drosophila suzukii*^11^, and *Aedes aegypti*^12^), the future prospects of RIDDLE are potentially promising. This is especially true considering that homing drives with good efficiency were also recently demonstrated in these species^38–40^. Indeed, because somatic expression and maternal deposition would not likely effect RIDDLE males, it should be substantially easier to build homing drives with high efficiency that are compatible with RIDDLE than to build high efficiency standard homing suppression drives.

Overall, we have shown that adding a homing drive to fsRIDL can substantially improve performance if resistance can be mitigated. Our proof-of-principle demonstration in *Drosophila* showed both high female lethality and high drive inheritance, suggesting the use of RIDDLE in major pest species. The self-limiting nature of this system may be particularly desirable in terms of securing regulatory approval, embodying beneficial characteristics from both powerful gene drive and well-established RIDL.

## Acknowledgements

This study was supported by laboratory startup funds from Peking University and the Center for Life Sciences, as well as the grants from the National Science Foundation of China (32302455, 32270672, and Overseas Youth). Thanks to the High-Performance Computing Platform of the Center for Life Science at Peking University for assistance with cluster-based data collection.

## Supplemental Information

### Supplemental Methods

**Figure S1.**
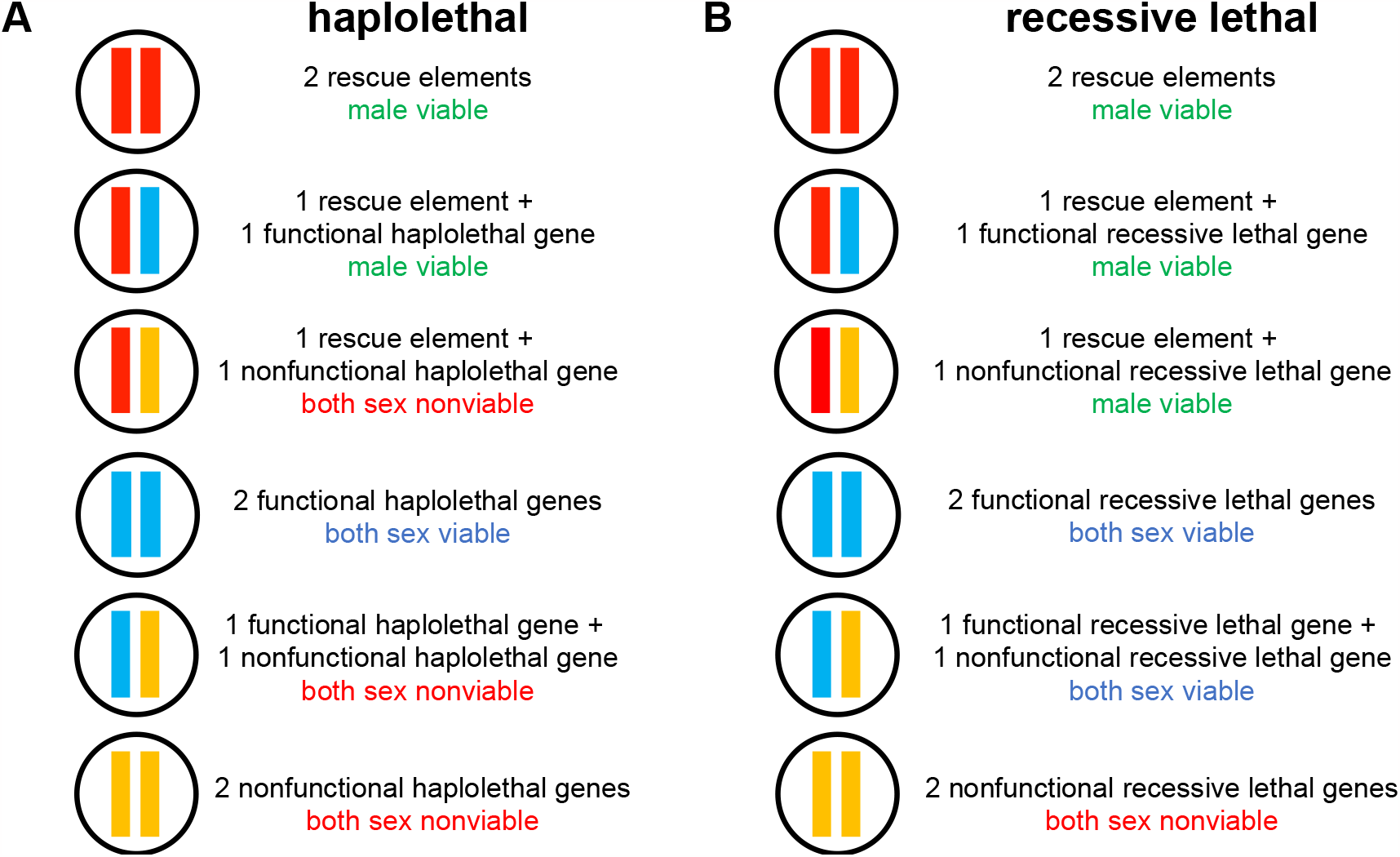
Rescue strategies of the RIDL-drive system. (**A**) Haplolethal drive. Individuals need two functional copies of the essential gene to survive, and therefore resistance carriers are nonviable. (**B**) Recessive lethal drive. Individuals need at least one functional copy of the essential gene to survive, and resistance homozygotes are nonviable.

**Figure S2.**
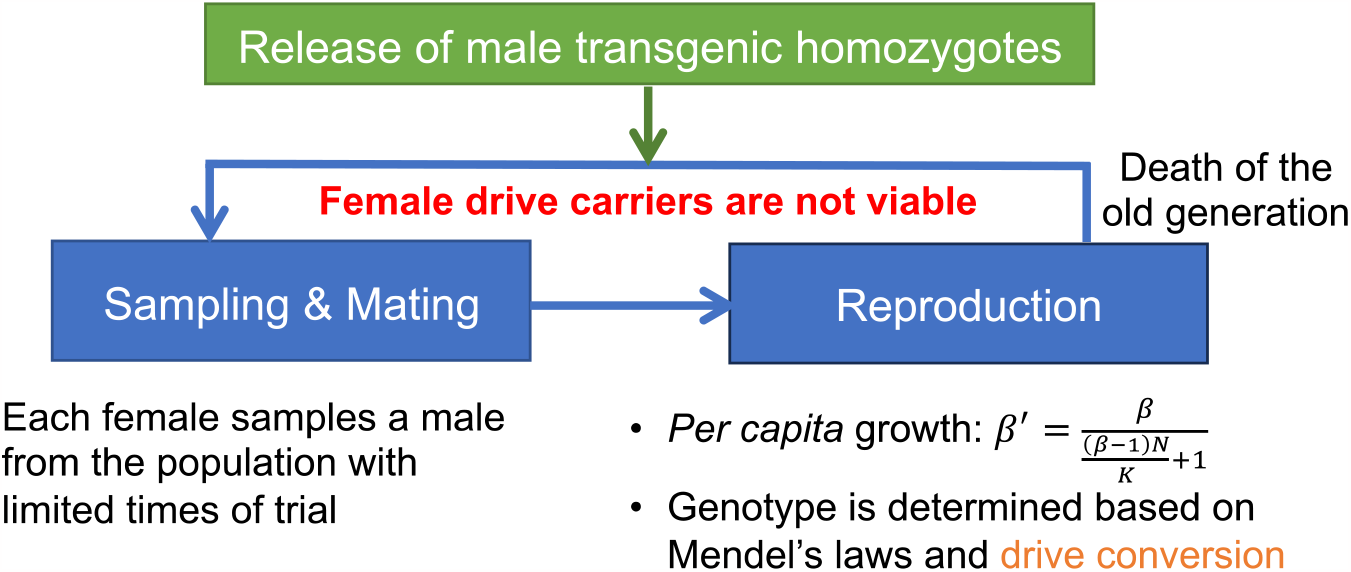
Workflow of the discrete generation model. The simulation is initiated by releasing males into a population. Each generation involves new reproduction based on density.

**Table S1.**
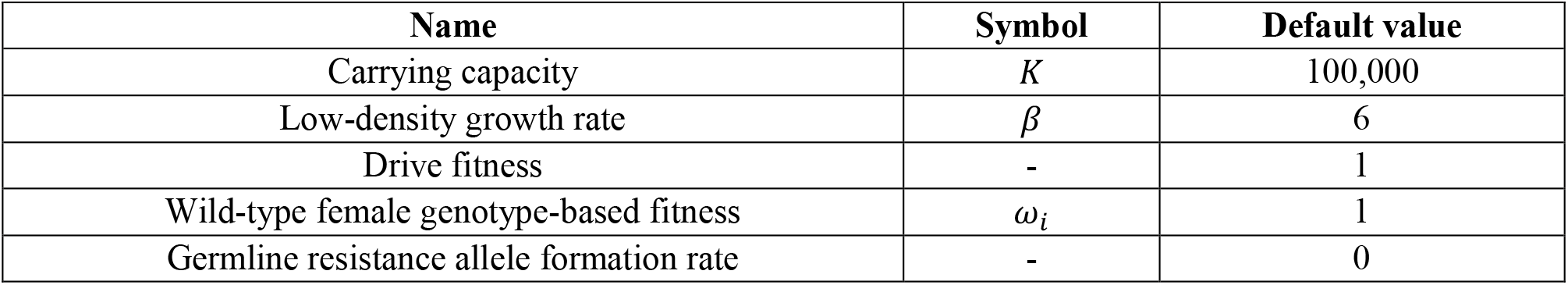
Default Parameters for the discrete generation panmictic model.

| Name | Symbol | Default value |
| --- | --- | --- |
| Carrying capacity | $K$ | 100,000 |
| Low-density growth rate | $\beta$ | 6 |
| Drive fitness | - | 1 |
| Wild-type female genotype-based fitness | $\omega_i$ | 1 |
| Germline resistance allele formation rate | - | 0 |

**Figure S3.**
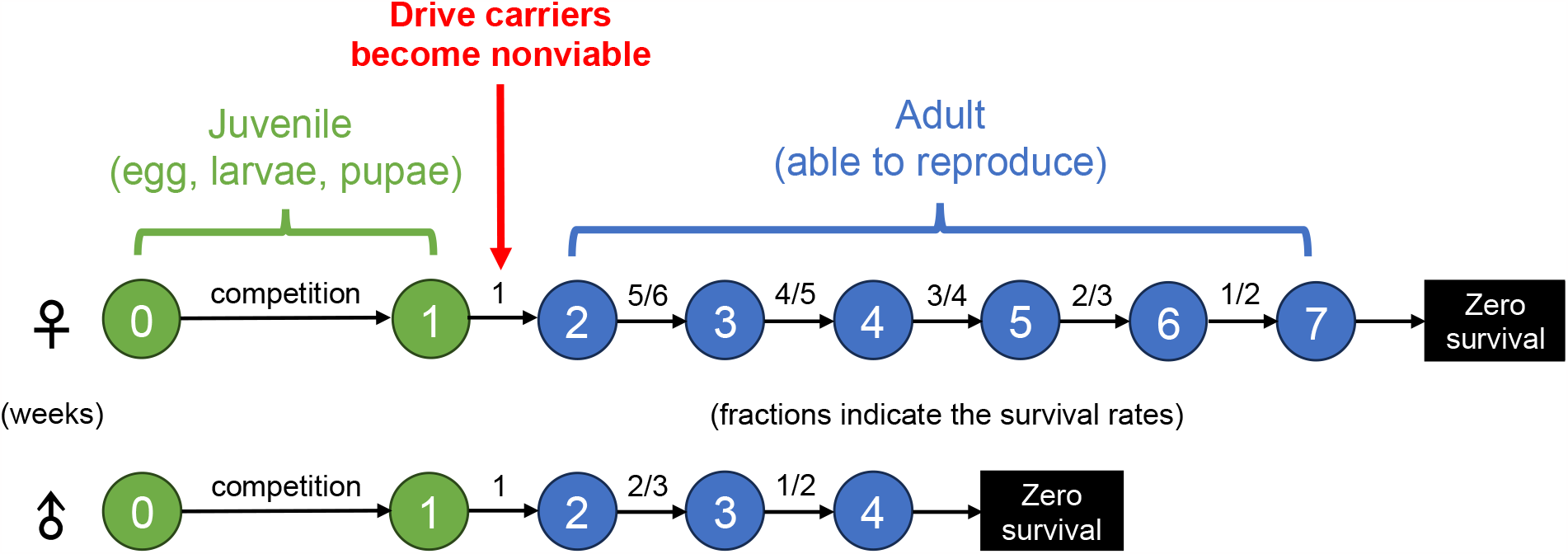
Workflow of the mosquito model. The simulation is initiated by releasing males into a population. Each weekly time step involves mating, new reproduction, density-dependent larval competition, and age-related mortality of adult mosquitoes.

**Table S2.** Default Parameters for the mosquito model.

| Name | Symbol | Default value |
| --- | --- | --- |
| Adult female capacity | $K_{\text{adult female}}$ | 50,000 |
| Low-density growth rate | $\beta$ | 6 |
| Expected number of offspring per adult female | $F$ | 25 |
| Drive fitness | - | 1 |
| Drive conversion rate (drive efficiency) | - | 0.5 |
| Germline resistance allele formation rate | - | 0 |
| Density-dependent growth curve | $\beta'(r)$ | Linear |

**Table S3.**
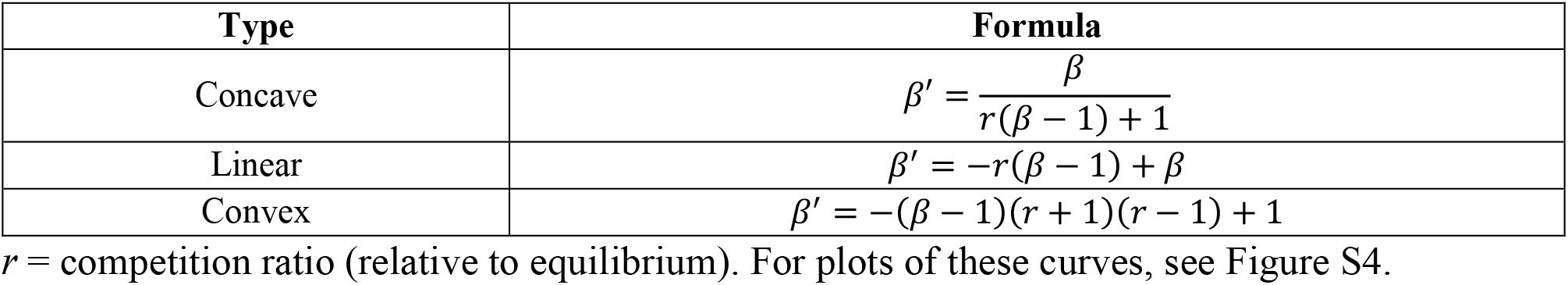
Different density-dependent growth curves (β*’*).

**Figure S4.**
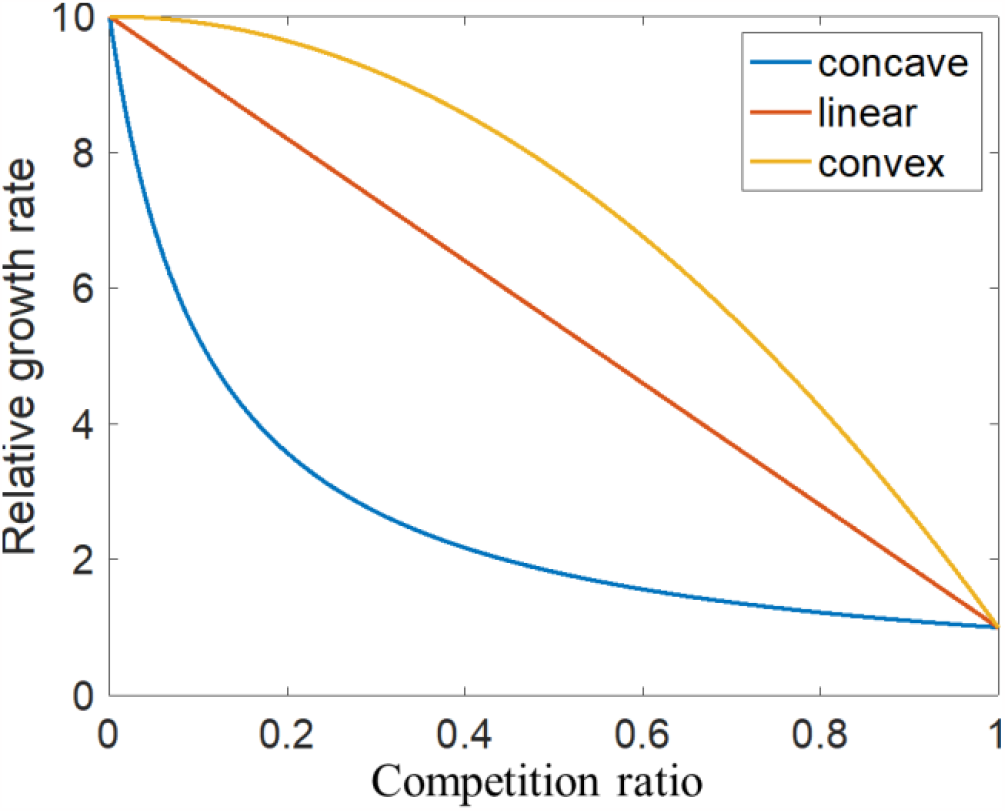
Density-dependent growth curves. The graph shows the relative survival advantage (or population growth rate) of larvae at lower population density (as defined by the competition ratio an individual experiences). For equations of these curves, see Table S3.

### Supplemental Results

**Figure S5.**
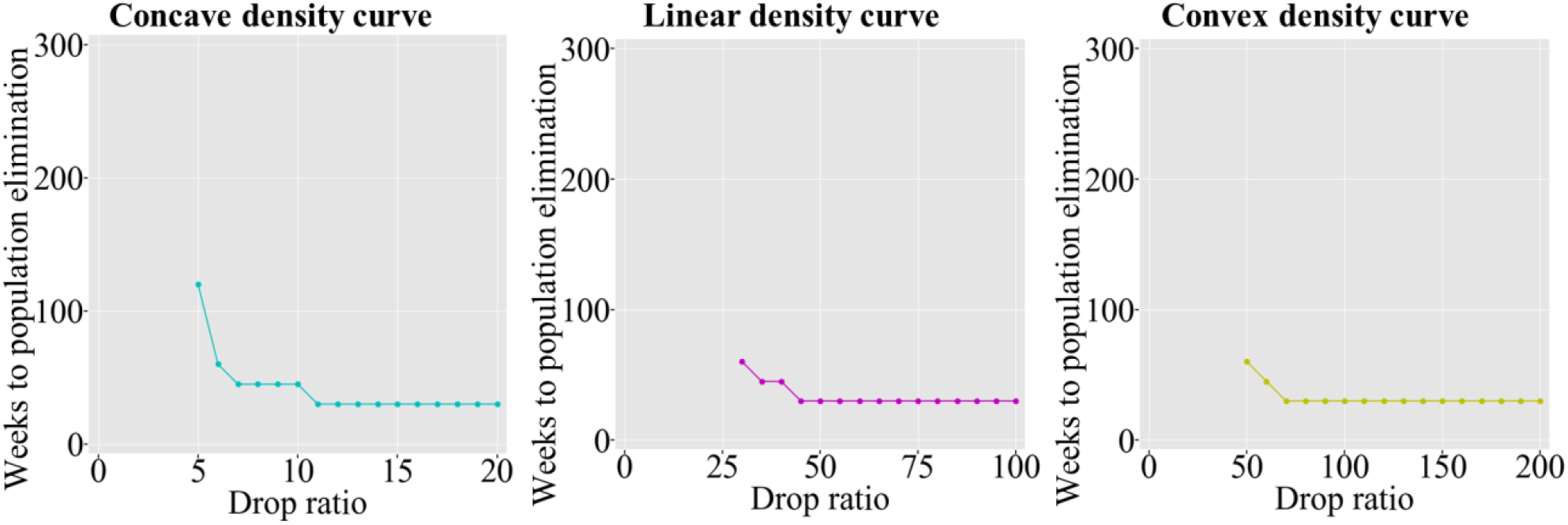
SIT performance with varying density dependence in the mosquito model. SIT males were released into a population every week based the drop ratio, which specifies the relative number released each generation (3.17 weeks) compared to the male population at equilibrium. The density dependence of the model was varied (see Figure S4), with a fixed low-density growth rate of 10. For low drop ratios, lack of a point indicates that population elimination did not occur.

**Figure S6.**
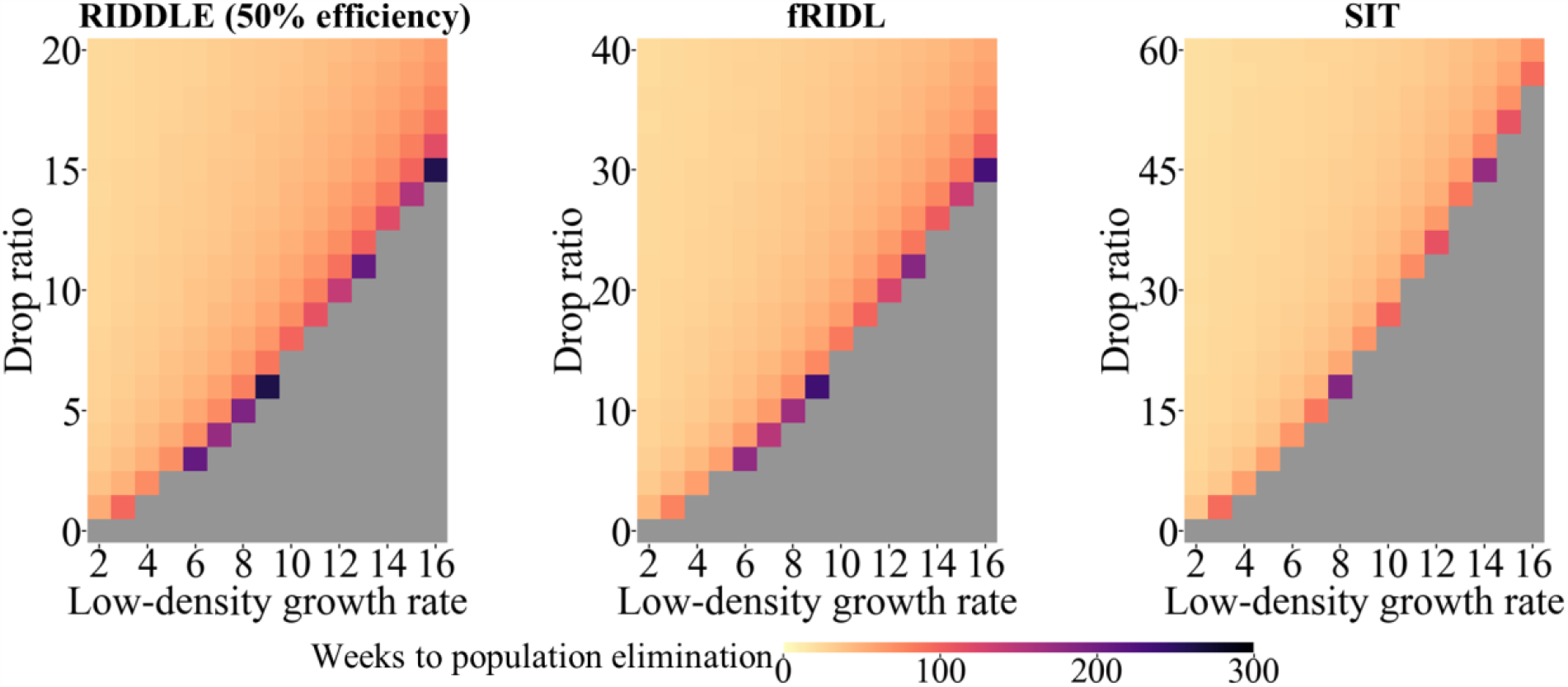
Effect of low-density growth rate in the mosquito model. Males of the indicated - types (RIDDLE males had 50% drive conversion efficiency) were released into a population every week based the drop ratio, which specifies the relative number released each generation (3.17 weeks) compared to the male population at equilibrium. The low-density growth rate varied, with a fixed low-density growth rate of 6 and linear density response curve. Grey indicates failure to eliminate the population after 100 generations (317 weeks).

**Figure S7.**
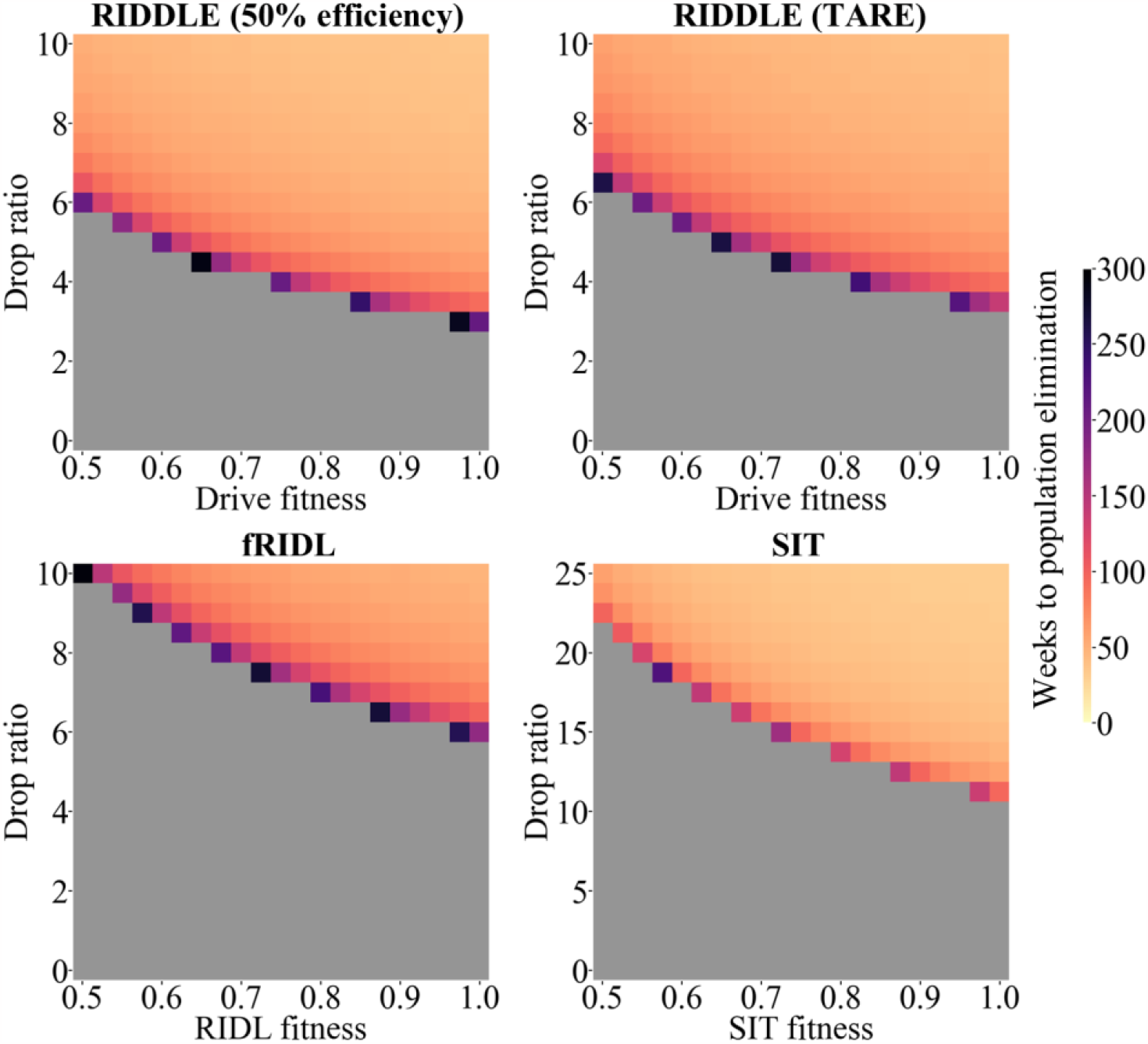
Effect of fitness costs in the mosquito model. Males of the indicated types (RIDDLE males had 50% drive conversion efficiency, and RIDDLE TARE males had a 100% germline cut rate)) were released into a population every week based the drop ratio, which specifies the relative number released each generation (3.17 weeks) compared to the male population at equilibrium. The fitness of each construct was varied, with a fixed low-density growth rate of 6 and linear density response curve. Grey indicates failure to eliminate the population after 100 generations (317 weeks).

**Figure S8.**
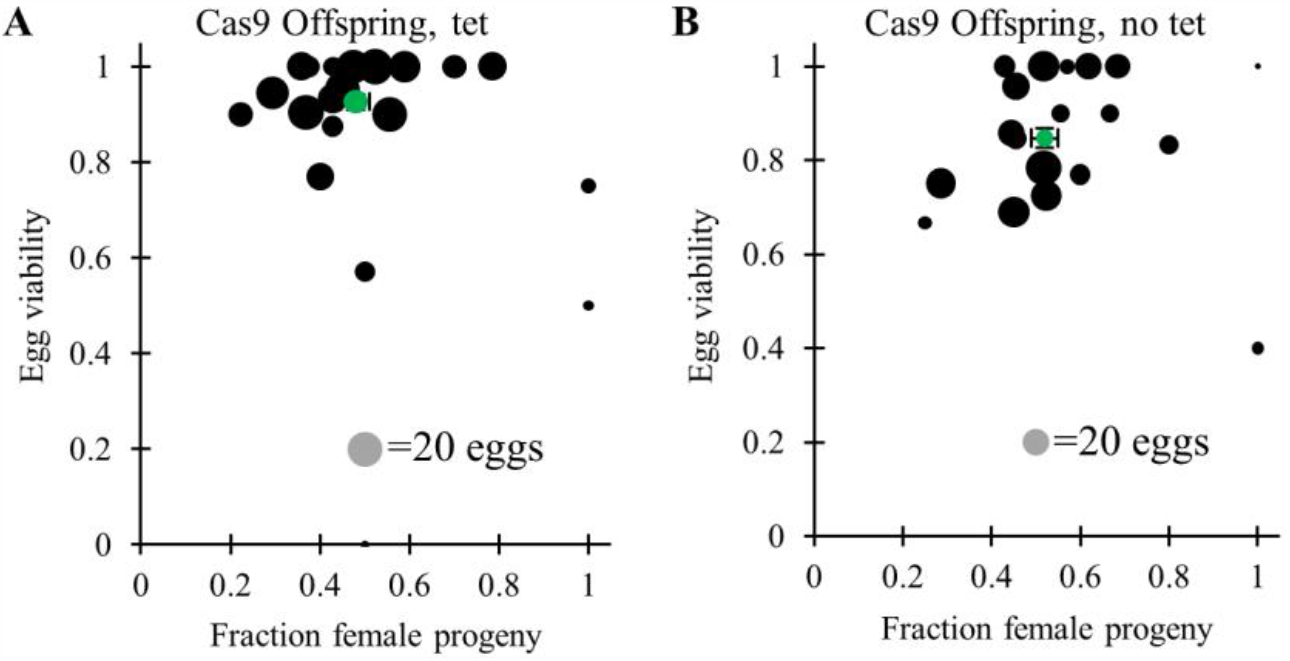
Egg viability and female fraction in control crosses with only Cas9. Several crosses with Cas9 homozygotes were set up, and egg viability was recorded in vials (**A**) with and (**B**) without tetracycline (tet). Adult progeny were also phenotyped for sex. Each dot represents progeny from a single drive individual. The green dot represents the mean for all individuals, and black bars represent the average and standard error.

**Figure S9.**
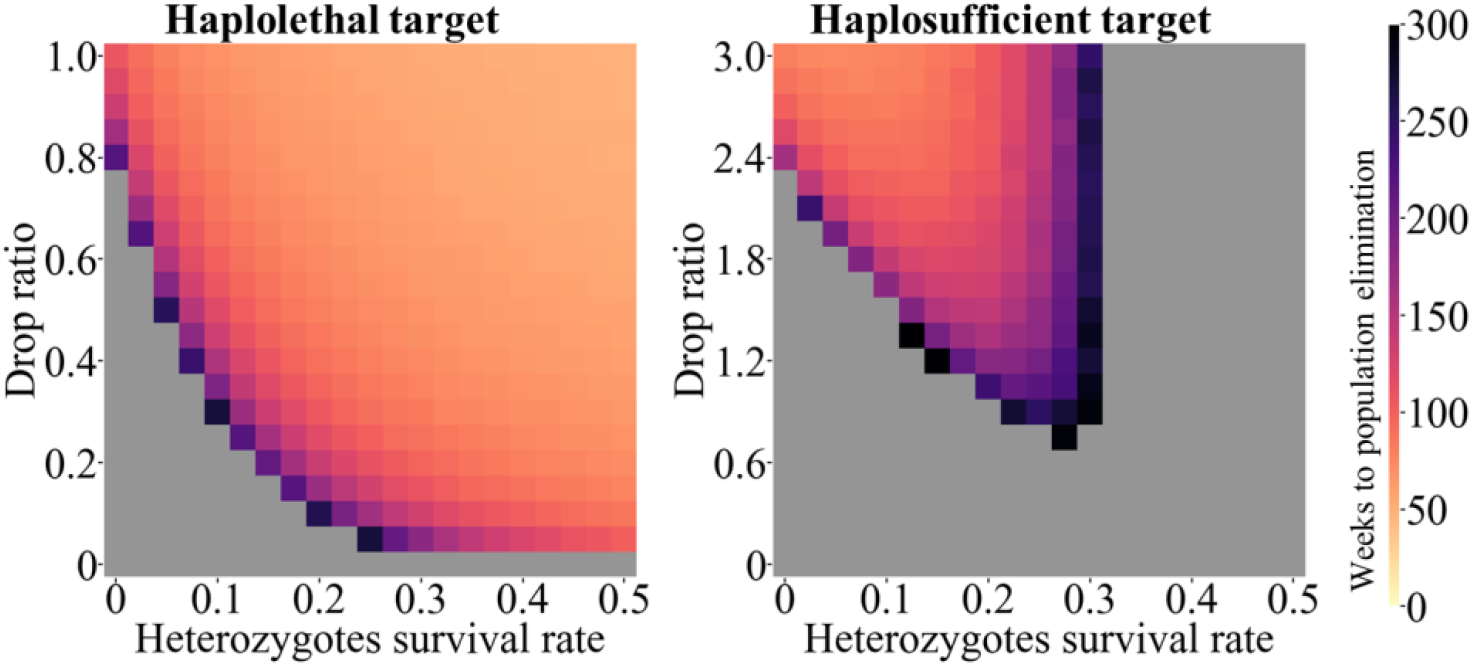
Effect of incomplete female heterozygote lethality. Males homozygous for RIDDLE were released into a population every week based the drop ratio, which specifies the relative number released each generation compared to the male population at equilibrium. With a low-density growth rate of 6, a drive conversion efficiency of 50%, and a nonfunctional resistance allele formation rate of 50%, the survival rate of heterozygous females was allowed to vary. Homozygote females were assumed to always be nonviable. Grey indicates failure to eliminate the population after 100 generations (317 weeks).

